# Auditory regularity detection in the ferret

**DOI:** 10.1101/2025.08.04.668417

**Authors:** Katarina C. Poole, Maria Chait, Jennifer K. Bizley

## Abstract

Acoustic sequences that transition from random to regularly repeating tones are increasingly used to study how the auditory system detects structure. Humans can identify such regularities rapidly, often within a single cycle. To test whether this ability is unique to humans, we trained ferrets (n = 6) to detect transitions from random tone sequences to repeating patterns. All animals learned the task and showed high accuracy for short (3 tone) patterns. Performance remained above chance for longer patterns (5 or 7 tones), though accuracy declined with increased pattern length. In a control condition where both random and regular segments contained the same five frequencies, ferrets continued to detect regularity, indicating they relied on temporal patterning rather than spectral content. To further rule out spectral cues, we included transitions from random 20 tone sequences to random 3, 5 or 7 tone sequences without repetition. Although these stimuli elicited performance above the false alarm rate seen in fully random trials, detection accuracy remained substantially lower than that observed for regular sequences. This suggests ferrets were not solely detecting changes in spectral statistics, but true regularity. Together, our results indicate that sensitivity to regular patterns within sound is not unique to humans and may reflect a broader auditory computation shared across species.

## 1. INTRODUCTION

The human brain continuously analyses the features of sounds within our acoustic environment to detect the emergence of new sources, extract their meaning, and adapt behaviour accordingly. Growing evidence suggests that the auditory system relies on statistical regularities to decompose acoustic scenes into these discrete objects (Denham and Winkler, 2006; Winkler et al., 2012, 2009). To investigate these mechanisms, studies often impose controlled regularities on sound sequences. One form is a deterministic repeating predictable pattern, such as a sequence of tones with consistent inter-tone-intervals and identically repeating frequency sequences. Alternatively, a pattern can be deterministic but aperiodic, where identical patterns are presented with individual elements within the pattern occurring at irregular intervals essentially producing a repeating rhythm, such as described by Asokan et al. (2021), or with regular spacing between repetitions, but irregular spacing within the pattern. In contrast to deterministic regularities, probabilistic regularities involve a degree of randomness rather than direct repetition. In this case, patterns emerge based on the likelihood of one sound following another. These probabilities can be learnt and range from simple rules, such as the mean and variance of frequencies in a tone cloud (Skerritt-Davis and Elhilali, 2018), to more complex statistical structures, like those governing language (Saffran et al., 1996). Regardless of whether regularity is deterministic or probabilistic, there is substantial evidence for the role of regularity in facilitating auditory stream segregation during auditory scene analysis (Andreou et al., 2011; Bendixen et al., 2010; Devergie et al., 2010; Rimmele et al., 2012; Southwell and Chait, 2018). Yet how the auditory system extracts and detects these statistical rules is still unclear.

Numerous studies have investigated how human listeners perform auditory pattern/regularity detection. Early work by Guttman and Julesz (1963) demonstrated that humans are sensitive to the repetition of broadband noise, even when the informational content is highly complex within noise. More recent research has refined this understanding by using pure tone sequences to parameterise the acoustic input more precisely. For instance, Barascud et al. (2016) illustrated the remarkable ability of human listeners to rapidly detect novel emerging patterns. In most cases, participants required less than one full repetition of a pattern to recognise the regularity (four tones after the initial presentation of the pattern). However, longer pattern lengths led to slower detection times, suggesting a strain on memory resources involved in tracking the repeated patterns. Differences in brain activation in response to the random to regular sequences vs. random sequences was observed using magnetoencephalography (MEG) during passive listening. These differences emerged at time points identical to that of the behavioural responses and to that of an ideal observer and indicated that regularity detection occurs automatically even when the stimulus is not behaviourally relevant (Barascud et al. 2016).

Research employing similar parametrically controlled stimuli to those in Barascud et al. (2016) have yielded comparable findings. For example, work by Southwell et al. (2017) and Herrmann and Johnsrude (2018) identified analogous sustained neural responses to regular tone sequences with electroencephalography (EEG). Additionally, Hu et al. (2024) found that these neural responses indicate regularity detection even with extended gaps between tones that effectively extended the pattern duration, albeit with a substantially reduced sustained neural response. Listeners exposed to the same regular sequence show performance advantages in detecting these patterns that persist for weeks or even months, suggesting that the patterns are stored in long term memory (Bianco et al., 2020; Herrmann et al., 2021). However, age-related differences in sensitivity to these patterns have been observed. Bianco et al. (2023) demonstrated that older listeners are slower to detect repeated auditory patterns and show a smaller performance advantages compared to younger individuals. Using a variable-order Markov model (Prediction by Partial Matching, or PPM model), Bianco et al. (2023) were able to simulate behavioural detection failures across age groups, linking these differences to variations in both early and long-term memory between younger and older listeners. Complementing the behavioural data, Herrmann et al. (2022) found that older adults exhibit reduced neural sensitivity to these repeated patterns compared to younger listeners. These results highlight how memory and cognitive processing capacities shape regularity detection, factors that may also help explain differences in pattern sensitivity observed across species. Nonetheless, despite these advances and a well-characterised understanding of how human listeners detect auditory regularities, the underlying neural mechanisms driving this ability remain poorly understood, highlighting the need for further investigation in this area.

One approach to elucidate the neural mechanisms is to use animal models. However, it remains unclear whether animals detect regularities in the same way as human listeners. Research has shown that animals are indeed sensitive to statistical regularities. For example, songbirds have been shown to learn and generalise dependencies between song elements (Chen and ten Cate, 2017; Petkov and Cate, 2020), while mice display a preference for regularity within mouse vocalisations (Perrodin et al., 2023). Neurophysiological recordings from animals during passive listening have also demonstrated a sensitivity to statistical patterns in auditory stimuli; Asokan et al. (2021) observed changes in neural firing rates depending on the regularity, with rhythmic (regular inter-stimulus interval) patterns eliciting more precise spike timing. Similarly Kang et al. (2021) reported a decrease in neural firing when a complex spectrotemporal pattern was repeated. Recent work by Bianco et al. (2024) showed sensitivity to regularity in timing, and less so in pitch, via pupil dilation and EEG in non-human primates. Although these neural responses indicate sensitivity to regularities, they only serve as proxies for perceptual processing. Behavioural evidence is necessary to determine the functional significance of these neural correlates. Moreover, investigating the neural mechanisms during active listening could offer additional insights into how statistical regularities are extracted.

In general, there are very few paradigms that show animals can detect deterministic regularities. One notable paradigm that uses regularity detection is figure-ground segregation, where the subject must detect a repeating pure tone chord (‘figure’) embedded in a mixture of random chords. Macaques are capable of performing this task (Schneider et al., 2021, 2018) albeit with a lower sensitivity to the figure than human listeners. A few studies have shown a non-primate model can detect statistical regularities: the ferret can identify the onset of regularly repeating single-frequency pure tone within a random tone cloud (Ma et al., 2010), suggesting they can segregate the repeating tone from the tone cloud. Ferrets are also able to detect a repeated frozen noise burst within a random noise mixture (Saderi et al., 2020). However, despite the complexity of the stimulus in the study by Saderi et al. (2020), only two unique target repetitions were used in each experimental block, making it unclear whether animals could detect the emergence of novel regularity on its first presentation.

To our knowledge, no animal model has been shown to detect transitions from random to regular tone sequences, as used by Chait and colleagues (Barascud et al., 2016; Bianco et al., 2020; Hu et al., 2024; Southwell et al., 2017). These tone sequences allow for precise control over the frequency content and length of the repeating patterns, enabling us to examine the auditory system’s ability to detect such transitions through behavioural tasks. Additionally, modelling the information content of these unfolding sequences using the PPM model provides a valuable benchmark for comparing human performance with animal performance, as well as grounding hypotheses about the cognitive mechanisms involved (Barascud et al., 2016; Bianco et al., 2023, 2020; Harrison et al., 2020). In this study, we selected the ferret (*Mustela putorius furo)* as our model organism firstly due to its hearing range which largely overlaps with human hearing (ferret range: approximately 20Hz to 44kHz; human range: approximately 20hz to 20kHz; Heffner and Heffner, 2007; Kavanagh and Kelly, 1988) and secondly their amenability to learning complex auditory tasks (Bizley et al., 2013; Keating et al., 2013; Ma et al., 2010; Nodal et al., 2008; Norris and Bizley, 2024; Saderi et al., 2020; Town et al., 2023; Town and Bizley, 2022; Walker et al., 2009; Yin et al., 2010) including ones that require elements of regularity detection and/or auditory stream formation (Griffiths et al., 2024; Lu et al., 2018; Ma et al., 2010; Saderi et al., 2020).

Using a paradigm extensively characterised in human studies, in this study we tested the ferret’s capacity to detect regularity. Specifically, we asked: can ferrets detect the transition from random to regular tone sequences; does their ability to detect regularity vary with different pattern lengths; what aspects of the stimulus structure facilitate or hinder detection; and do they adopt behavioural strategies similar to those observed in human listeners? By addressing these questions, our study aims to determine the extent to which ferrets can serve as a viable model for investigating the neural basis of auditory regularity detection. Our findings reveal that ferrets can indeed detect transitions from random to regular tone sequences. We report that they show improved detection with simpler or more predictable patterns and exhibit behavioural signatures consistent with those of human listeners. These results position the ferret as a valuable animal model for probing the mechanisms underlying regularity detection and auditory scene analysis.

## 2. METHODS

All experimental procedures performed were first approved by a local ethical review committee. Procedures were carried out under license from the UK Home Office in accordance with the Animals (Scientific Procedures) Act (1986) and PPL: PP1253968.

### 2.1 Animals

Six adult female-pigmented ferrets (*Mustela Putorius furo*, Highgate Farm, UK) were trained in the regularity detection task (F1805, F1812, F1813, F2001, F2003, and F2103). Ferrets were housed in groups of two to eight in enriched cages. All had regular otoscopic examinations to make sure ears were healthy. Animals were water regulated for behavioural testing. During regulation, which typically lasted from Sunday afternoon until Friday afternoon, subjects were weighed daily to ensure that their body weight did not drop below 88% of their initial weight. A minimum of 60 ml/Kg was provided to the animal daily either through water rewards during the task, water manually supplemented by the experimenter, and/or mash (crushed food pellets mixed with water) provided to each water restricted ferret at the end of the day.

### 2.2 Stimuli

Stimuli were generated in MATLAB and presented at a sampling frequency of 24414Hz. All frequencies were individually level calibrated at the position of the animal’s head using a microphone (Brüel & Kjær ½” Microphone 4134), measuring amplifier (Brüel & Kjær 2610) and tone generator (Brüel & Kjær 4231), between a noise-floor of 30 to 40 dBSPL to maximum presentation of 70 dBSPL to minimise non-linearities in frequency presentation within the speaker.

#### 2.2.1 Regularity detection

Acoustic stimuli consisted of sequences of 50 ms tone pips (with 5 ms cosine on/off ramps), presented with no inter-tone interval, drawn from a pool of 20 that comprised 120 Hz to 9.7 kHz with third octave spacing. This was identical to that of the studies performed by the Chait lab but with a single adaptation (third octave spacing rather than sixth) to account for the wider auditory filters in ferrets than that of humans (Sumner et al., 2018). From this pool, tones were randomly drawn to create a random tone sequence (RAN) which on 50% of trials transition to a regular tone sequence (RAN-REG) 1.5 to 2.5 s after stimulus onset. Regular sequences were generated by selecting *n* tones (where *n* signifies the pattern length that could be 3, 5 or 7) from the transition time to then be repeated, with the regular sequence lasting 2 s (see Figure 1A). The transition time was randomised for each trial between 1.5 to 2.5 s (in 50 ms steps, uniformly distributed) after stimulus onset. These entire sequences were randomly attenuated between the ranges of 54 to 60 dBSPL at 1.5 dB steps. Each tone sequence was newly generated for every trial apart from a subset of sessions where sequences were repeated up to 10 times within a session to provide reliable spiking responses for electrophysiological recordings in later experiments. In the repeated presentations the transition time was fixed at 2 s and the level at 60 dBSPL. RAN trials comprised of randomly repeating tones with a sequence duration of 3.5-4.5s to match the duration of the RAN-REG trials (see “Training and testing” below).

**Figure 1:**
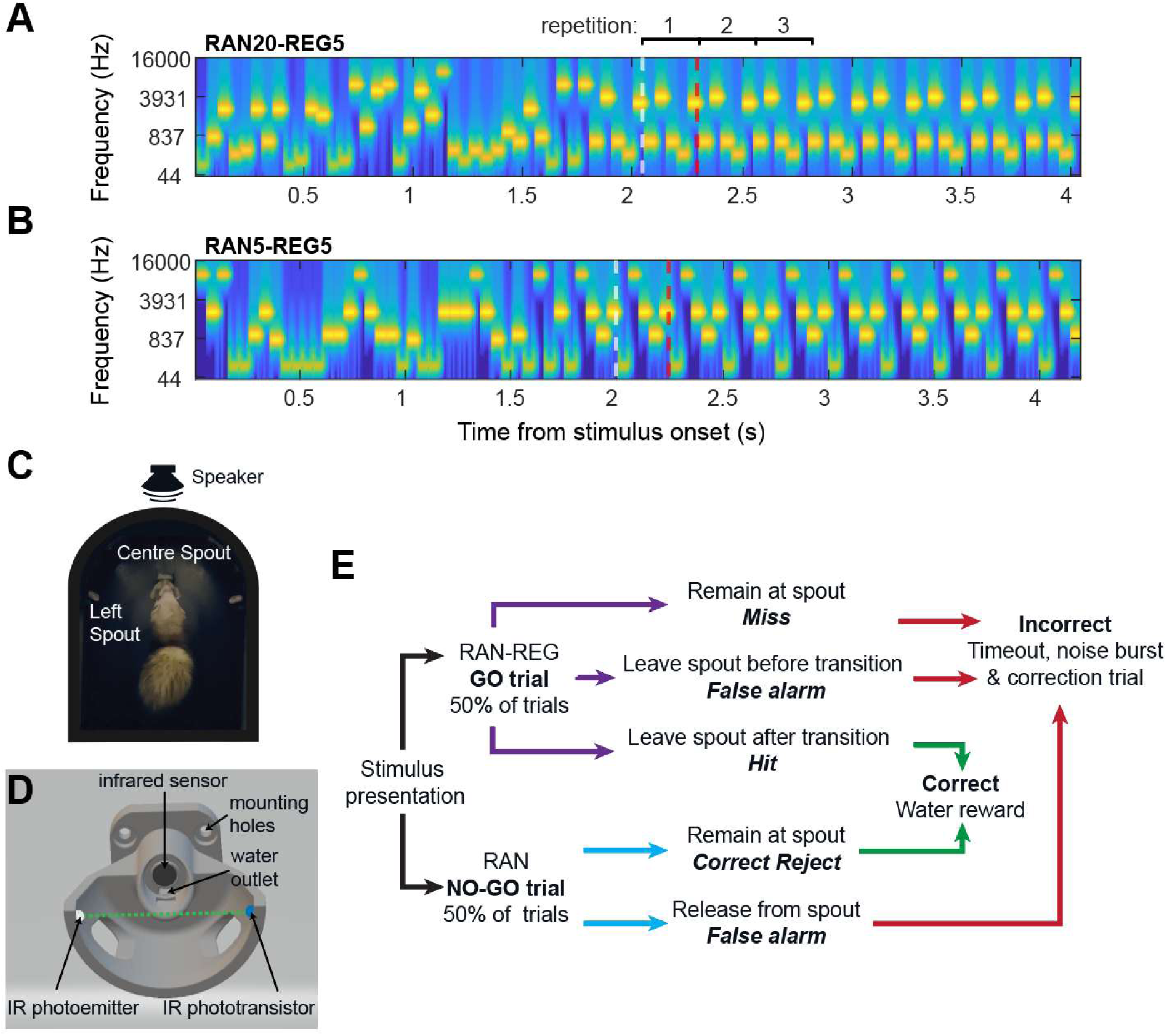
Experimental stimuli, apparatus and procedure. A-B) Sequences consisted of 50 ms tone pips and ranged in length from 3.5 to 4.5s A) Example random sequence drawn from an alphabet of 20, that transitions to a regularly repeating pattern of 5 tones (RAN20-REG5). White dashed line indicates the transition time, red dashed line shows the end of the cycle. B) Example of a sequence random alphabet of 5 frequencies, that transitions to a repeating pattern of 5 tones (RAN5-REG5). C) Behavioural testing was housed in a double-walled sound attenuated booth. D) Performance was monitored with a custom 3D printed structure that housed an infrared beam break (green dashed line) and infrared reflecting sensor (black circle) to monitor behaviour and a water outlet to provide a water reward. E) Schematic of trial structure for both fully random (RAN, NO-GO trials) and trials that transition to a regular sequence (RAN-REG, GO trials).

#### 2.2.2 Control tasks

To probe whether the animal uses the deterministic pattern during detection to perform the task, rather than other cues such as changes in spectral content, two further conditions were tested. Firstly, a condition where the random tone sequence only consisted of five randomly chosen frequencies (RAN5) that were followed by a pattern of 5 tones (REG5) using the same preceding frequencies (see Figure 1B). This prevented animals from using any information about the probability of occurrence of any one tone as a cue for detection. The second was a random sequence (RAN20) that transitioned to a random sequence of 3, 5 or 7 tones (RAN20-RAN3, RAN20-RAN5, RAN20-RAN7 respectively; see Figure 4A). This condition captured the change in spectral statistics that occurred in the RAN20-REG*n* conditions, without introducing a regular pattern. These trials were introduced as probes on a maximum of 10% trials within a session and were randomly rewarded to avoid biasing trained behaviour.

### 2.3 Apparatus

Animals were trained in a double-walled sound attenuated booth in a semi-circular arena encased in acoustically transparent plastic mesh (see Figure 1C). The stimulus was presented from a free field central loudspeaker (Visaton SC 5.9) approximately 20cm from the subject’s head position. A central spout on a post in the centre of the arena held an infrared (IR) sensor (Optek / TT Electronics OPB710) and IR beam breaker consisting of an emitter (Optek / TT Electronics OP165A) and phototransistor (Optek / TT Electronics OP506A) contained in a custom 3D printed structure (see Figure 1D and Poole (2025) for the 3D spout models). The central spout also delivered water, controlled by a solenoid (Valeader B2B112.BVO 12VDC) to provide water rewards (approximately 0.25 ml per reward). A secondary spout to the left-hand side of the central post (approximately 15 cm away) was used as the response location for when the animal was required to respond after the transition from the random to regular sequence. This spout contained just an infrared sensor and a channel for the water reward. The experiment was run on custom scripts written in MATLAB (The Mathworks) which communicated to the TDT signal processor (RX8) to control stimulus presentation and receive sensor information. One camera (Intel Realsense D435) was installed to observe the animal during the experiment.

### 2.4 Training and testing

Fully trained animals completed the task with RAN-REG sequences presented on 50% of trials, with the transition occurring between 1.5 to 2.5 s (see Figure 1E for trial structure). Animals were rewarded for releasing from the centre spout within 2 s of the start of the REG sequence. RAN trials were catch trials in which only random tones were presented and were 3.5 to 4.5 s in duration. Animals were rewarded at the end of the sequence from the centre spout if they had maintained contact throughout the full stimulus period. Early releases, releases during RAN trials and misses were punished with a noise burst and a brief timeout.

To train animals, ferrets were firstly positively reinforced to associate water with both the central and left spouts. Acoustic discrimination began with very short NO-GO trials and long regular (GO) tone sequences. As the subject’s performance increased after multiple training sessions the time between the stimulus onset and transition was gradually increased up until the testing conditions of the range 1.5 to 2.5 s; thereby the response window was gradually decreased from 10 s to 2 s. Once the animal was able to successfully detect the change from random to regular in the RAN20-REG3 condition within the 2s response window, increasingly longer pattern lengths (5 then 7 tones) were then introduced. These were initially presented in separate sessions but then all pattern lengths were randomly interleaved within a session. The RAN5-REG5 condition was presented separately and only after the animal became competent detecting the other conditions. Animals were considered trained when their performance was stable over time for each of the three pattern lengths. Sessions prior to this point were excluded from analysis. Probe trials (RAN20-RANX, where X = 3, 5 or 7) were also presented on a subset of sessions, with probes occurring randomly in <10% of trials. Probe trials were randomly rewarded with 50% probability.

### 2.5 Behavioural analysis

All behavioural data extraction and analysis was performed offline using Python and MATLAB. A response was a hit if the trial was a GO trial and the animal responded at the left spout after the transition time and before the end of the response window (2s) and a miss if they remained at the centre spout (see Figure 1E). A correct reject was logged if the animal held at the centre spout during a NO-GO trial. False alarms were logged when the animal release from the centre before the transition time on a GO trial or at any time during a NO-GO trial. Percent correct was calculated for each session as follows:

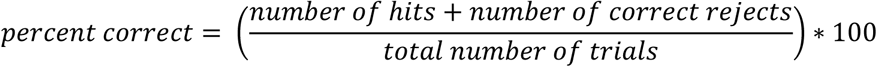

Sessions were excluded from further analysis if the animal scored below 40% correct where chance performance was 50%. On average, 25.8% of sessions per ferret were removed for this reason, most often on Monday mornings after water regulation began the previous day or on days when equipment failed. We also excluded any session in which the false alarm rate exceeded 65%, which affected on average 4.54% sessions per ferret.

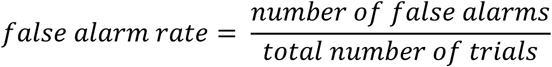

The first four trials of every session and trials in which the animal was registered as releasing from the centre within 500 ms of trial initiation were excluded from analysis (the later usually indicating failures to properly initiate the trial). To assess reaction time, sessions where the ferret performed at greater than 60% correct only were included to avoid analysis of chance responses.

To examine whether each ferret performed above chance in each condition, we used a permutation test to build a null distribution of detection performance. For each animal and each condition separately, we took the full set of trial-level responses, which includes whether the ferret withdrew from the central spout and the time of that withdrawal, and randomly shuffled those responses across trials. This preserves the overall response frequency and timing but removes the link between the response and original trial type. On each shuffled dataset, we then recomputed hit and false alarm rates such that any shuffled withdrawal on a NO-GO trial counted as a false alarm, and any shuffled withdrawal on a GO trial whose shuffled response time fell after the sequence transition counted as a hit. This process was iterated 1,000 times per ferret and condition, yielding a distribution of chance-level performance. Finally, we took the 97.5th percentile of that distribution as the threshold for classifying a ferret’s observed performance as significantly above chance.

Statistical analysis of effects of pattern length and effects of matched/unmatched alphabets and other conditions used general linear models and generalized linear mixed models fitted using fitglme in MATLAB (version 2022a). The details of each model are outlined alongside the relevant results; however, in general, analysis of behavioural performance (correct vs. incorrect responses) was based on a logistic regression in which the generalised linear model used a binomial distribution and logit link function settings and analysis of reaction times was based on a general linear model. For each model, the ferret was used as a random factor and we reported the magnitude of coefficients (estimate) of fixed effects of interest, the t-statistic for a hypothesis test that the coefficient is equal to 0 (T) and its respective p-value (p). The 95% confidence intervals are also reported for each fixed-effect coefficient and the adjusted R2 value of the model to assess model fit.

### 2.6 PPM and PPM-Decay modelling

To gain insight into how ferret behaviour differed from human behaviour in the same task, we employed both the standard Prediction by Partial Matching (PPM) model and a modified version, the PPM-Decay model. The PPM model is a variable-order Markov model that predicts upcoming events by combining statistics from n-gram models of varying orders (Cleary and Witten, 1984), using interpolated smoothing to balance the specificity of high-order patterns with the stability of lower-order ones (Bunton, 1997; see for more details: Harrison et al., 2020; Pearce and Wiggins, 2004). In previous cognitive applications (Barascud et al., 2016; Cheung et al., 2019; Gold et al., 2019), the PPM model assumes perfect memory, treating all n-gram observations with equal weight, regardless of recency. However, human memory shows clear capacity and recency effects. To reflect this, Harrison et al. (2020) introduced PPM-Decay, in which memory fidelity decreases over time via a customisable decay function. This model includes a high-fidelity echoic memory phase followed by a short-term memory stage with exponential decay.

In the present study, we employed both the standard PPM and the PPM-Decay models to simulate how an ideal observer, with perfect memory, and a memory-constrained observer might respond to transitions from the random to regular sequences presented. This approach aimed to account for the differences observed between human and ferret performance. For each stimulus condition (e.g. RAN20-REG3, RAN20-REG5, etc.) we generated 1000 simulated trials with a fixed transition point at 2 s. Frequencies were mapped to integer indices (1 to 20), and both models were initialised with an alphabet size of 20. To ensure unbiased learning of pattern emergence, the model was reinitialised on each iteration. In the PPM-Decay model, memory constraints were imposed using an arbitrary long-term memory half-life value of 0.09. For both models, we extracted the information content of each tone in the sequence which may be used by the auditory system to detect transitions from random to regular sequences.

## 3. RESULTS

### 3.1 Ferrets exhibit robust regularity detection and their behaviour is influenced by pattern length and alphabet size

Ferrets (n = 6) were trained on a GO/NO-GO-task where they were required to respond at a peripheral spout when they heard a transition from a random sequence to a regular sequence. The NO-GO trials were random sequences of frequencies either selected from a pool of 20 (RAN20) or a pool of 5 (RAN5). GO trials were random to regular sequences that either went from a pool of 20 frequencies down to a repeating pattern of 3, 5 or 7 tones (RAN20-REG3, RAN20-REG5 and RAN20-REG7 respectively); or a pool of 5 frequencies down to a repeating pattern of 5 tones (RAN5-REG5). All ferrets successfully performed the task for all tested conditions (defined as performing greater than the 97.5th percentile of the randomly permuted chance distribution; range of 97.5th criterion across ferrets = 47.475 to 55.312, as seen in Figure 2A-B). Mean performance (% correct) for each ferret in each condition can be seen in Table 1.

**Figure 2:**
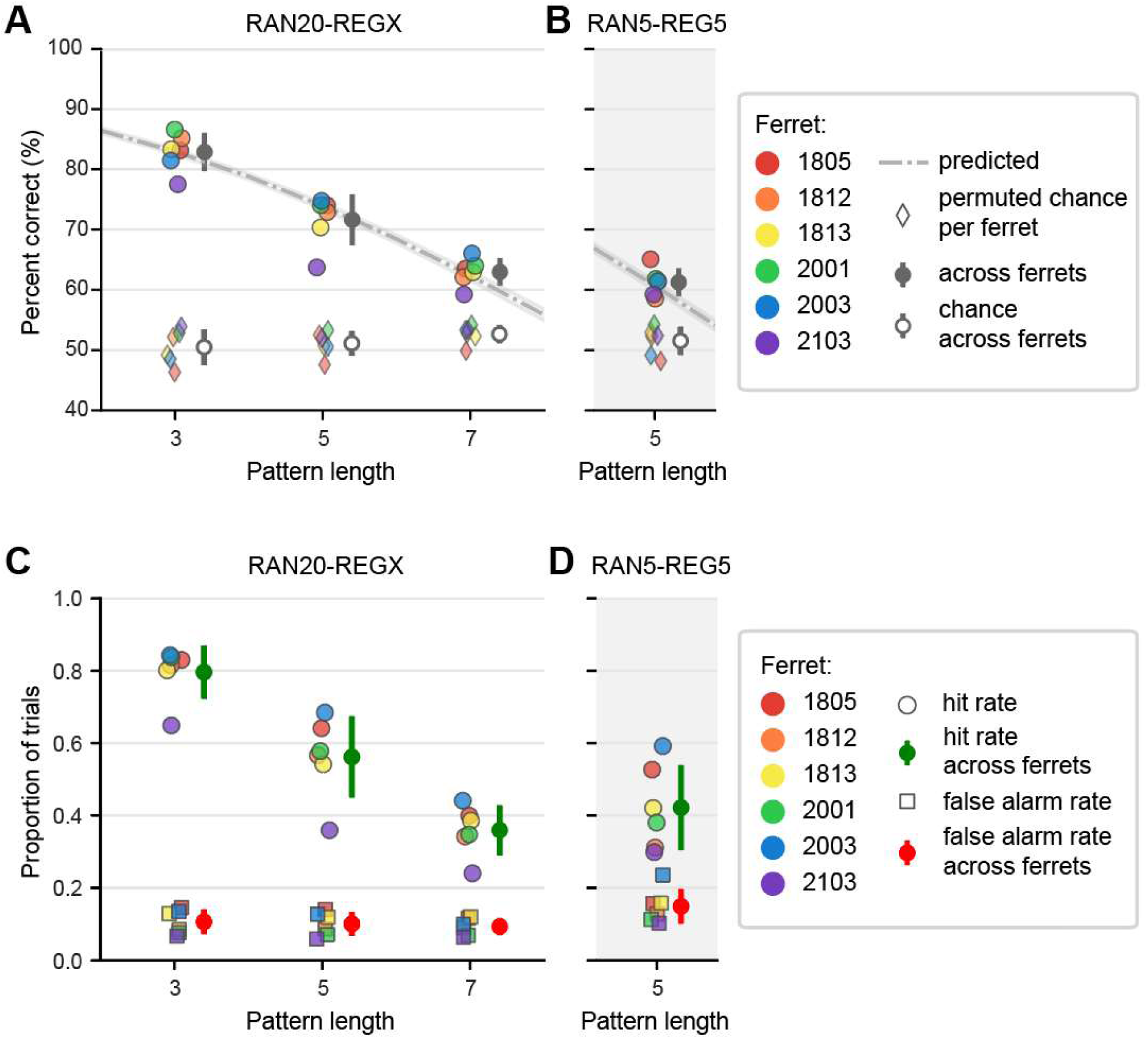
Performance of ferrets to increasing pattern lengths and random alphabet sizes. A-B) Average performance for each ferret (coloured circles) with the mean (±standard deviation) across ferrets shown in grey for (A) RAN20 and (B) RAN5 trials. Square markers indicate chance performance (calculated with a permutation test, see methods). C-D) Hit rate on GO trials (circle) and false alarms (square) across trials are displayed for each ferret and condition. Overall hit rate (green) and false alarm rate (red) is calculated across ferrets for the RAN20 (A) and RAN5 (D) conditions.

**Table 1:** Mean and standard deviation of performance (percent correct) of each ferret in each condition and the number of total trials in brackets.

| Ferret | RAN20-REG3 | RAN20-REG5 | RAN20-REG7 | RAN5-REG5 |
| --- | --- | --- | --- | --- |
| **F1805** | 83.14 ± 12.43 (5762) | 74.02 ± 13.70 (2236) | 63.51 ± 15.56 (766) | 65.08 ± 6.90 (3637) |
| **F1812** | 85.18 ± 10.75 (9797) | 72.91 ± 11.86 (6885) | 62.09 ± 12.97 (4313) | 58.54 ± 5.75 (7134) |
| **F1813** | 83.33 ± 10.96 (8410) | 70.35 ± 12.86 (5664) | 62.99 ± 13.72 (2626) | 61.39 ± 8.13 (2802) |
| **F2001** | 86.59 ± 8.73 (9632) | 74.06 ± 10.19 (7178) | 63.98 ± 10.48 (4249) | 61.83 ± 5.13 (8754) |
| **F2003** | 81.46 ± 9.95 (8740) | 74.77 ± 11.13 (8231) | 66.03 ± 9.53 (3902) | 61.44 ± 5.87 (5205) |
| **F2103** | 77.53 ± 8.66 (3098) | 63.71 ± 8.50 (3055) | 59.24 ± 10.30 (2211) | 59.24 ± 5.39 (2868) |

Having established that all ferrets are able to perform the behavioural task above chance levels for all conditions tested, we wanted to ask how the length of the pattern in the regular sequence affects behavioural performance. As can be seen in Figure 2A-B, performance for detecting regularity within the RAN20 condition appears best for the shortest pattern lengths. It is also apparent that the RAN5 condition elicited worse performance than trials with the same pattern length but drawn from an alphabet of 20 (RAN20-REG5). This was confirmed with a binomial mixed effects regression model (see Supplementary Table 2) that shows a main effect of pattern length, decreasing performance as the pattern length increases (β = −0.270, p < 0.001) and a main effect of the random alphabet, increasing performance with increasing alphabet size (β = 0.0406, p < 0.001). These findings are further supported by examining changes in hit rate and false alarm rate across conditions (see Figure 2C-D). False alarm rate remained relatively stable at approximately 10% across all conditions, indicating a consistent baseline response. In contrast, hit rate declined with increasing pattern length, demonstrating reduced sensitivity to longer patterns. Overall, this confirms that ferrets are better able to detect shorter patterns, and benefit from the change in spectral probabilities that accompanies the change from RAN20-REGX.

We then examined reaction times as a function of pattern length. Previous studies have shown that for patterns containing up to 10 tones, human participants typically require one full cycle of the pattern plus four additional tones to respond, consistent with predictions from an ideal observer model. Reaction times in humans then increase for patterns longer than 10 tones, an effect thought to reflect limitations in memory capacity (Barascud et al., 2016). Trained ferrets also showed an increase in mean reaction time for longer patterns (see Figure 3A, C). We ran a mixed effects linear regression (see Supplementary Table **5**) that confirmed a main effect of the pattern length, increasing reaction times with increasing pattern lengths (β = 0.0461, p < 0.001), such that an increase in the pattern by one tone increases the reaction time by approximately 46 ms. The model also indicated a main effect of the random alphabet, increasing reaction times with increasing alphabet size (β = 0.00503, p < 0.001), revealing shorter reaction times for RAN5-REG5 than for the RAN20-REG5 condition (see Figure 3B, D).

**Figure 3:**
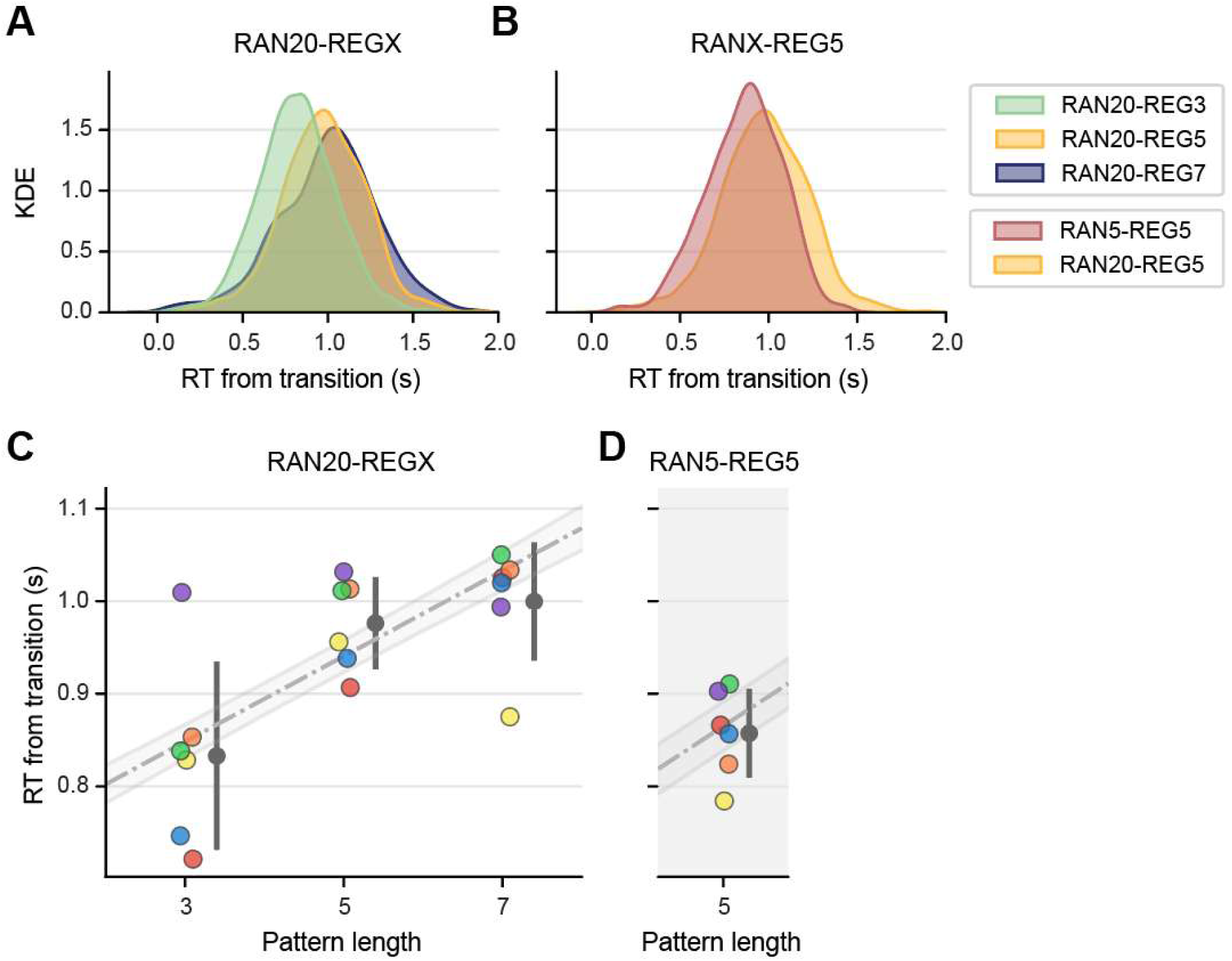
Reaction times of ferrets to increasing pattern lengths and random alphabet sizes. A-B) Kernel density estimates of reaction times, pooled across ferrets, to GO trials in sessions with > 60% correct separated by pattern length for RA20 condition (A) and separated by random alphabet for the PL5 condition (B). C-D) Average reaction time for each ferret and condition with the predicted values from the mixed effects linear regression shown (grey dashed line).

### 3.2 Ferrets detect the repeating pattern rather than a change in stimulus statistics

The analysis above provides evidence that ferrets are able to detect transitions from random to regular sequences. However, this task could potentially be solved by detecting the change in spectral statistics associated with the transition from 20 possible frequencies down to 3, 5 or 7, frequencies that constitute the pattern. While subjects were able to detect the emergence of a pattern without a change in spectral statistics in the RAN5-REG5, performance was significantly worse compared to the unmatched counterpart (RAN20-REG5). To understand to what extent ferrets were monitoring the spectral frequency probability, we designed probe trials in which there was a change in the number of frequencies at the transition, but no pattern (RAN20-RANX, see Figure 4A). These were presented as probe trials on <10% of trials within a session and randomly rewarded.

**Figure 4:**
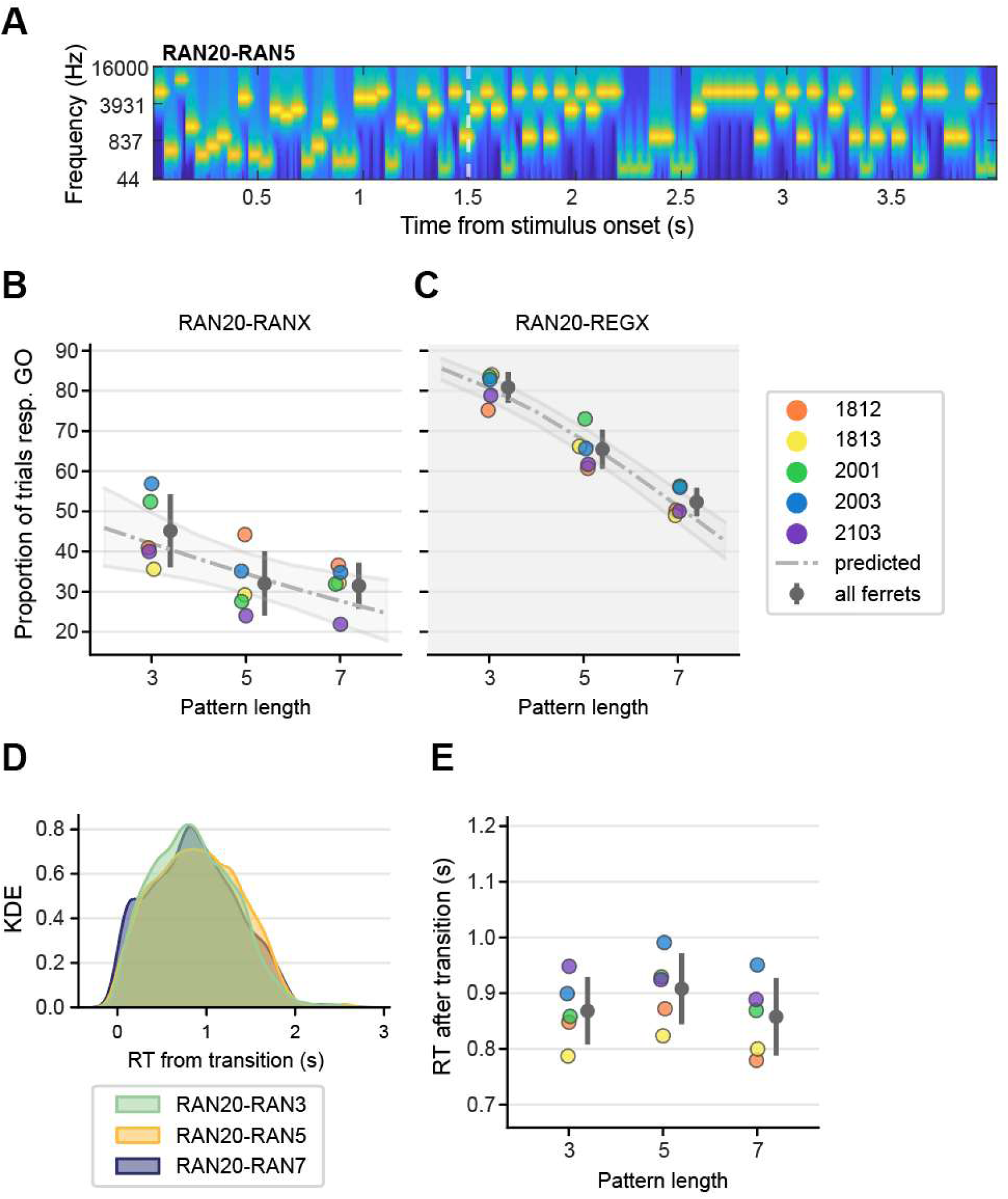
Performance and reaction time on probe trials with a change in stimulus statistics but no repeating pattern. A) Spectrogram of an example probe trial in which a random sequence drawn from 20 frequencies, transitions (at the dashed white line) to a random sequence drawn from 5 frequencies. B) Proportion of probe trials in which the ferret responded to the left (GO response). B) Proportion of trials containing a pattern (GO trials) in the same sessions in which the subject made a GO response. C) Kernel density estimate of all reaction times on probe trials across all ferrets, split by alphabet size. D) Reaction time from transition from random alphabet of 20 to 3, 5 and 7 for each ferret. Error bars = std. Model predictions in grey dashed line with 95% confidence intervals.

Figure 4B plots the proportion of trials in which ferrets responded GO on the RAN20-RANX probe trials; hit rates were much lower than when they heard the repeating pattern (see Figure 4C) although higher than their general false alarm rate (which is around ∼10%; see Figure 2C-D), particularly for the largest changes in statistics (RAN20-RAN3). A mixed effects binomial regression (see Supplementary Table **4**) revealed main effects of probe vs. pattern, and pattern length as well as an interaction between these factors. To determine the effect of pattern length on probe trial individually and to compare these to pattern trials, separate models were run on trials containing the probe and trials without. This revealed a smaller coefficient for pattern length in the probe condition (β = −0.154, p = 0.00459) compared to that of the pattern conditions (β = −0.334, p < 0.001). Therefore, if the stimulus just contained a drop in statistics and without a pattern the animal is significantly less likely to respond GO, suggesting that the ferrets are genuinely using the repeating pattern as marker to respond GO in the task.

We compared reaction times for probe and pattern stimuli (see Figure 4C-D), observing that for probe trials there was no modulation of reaction time by ‘pattern’ length, and that on average reaction times were longer for probe trials than pattern trials. This was confirmed with a mixed linear regression that showed no effect of pattern length (p = 0.861) but did show a main effect of whether there was a pattern or a probe, where if it was a probe trial the reaction time significantly increased compared to when a regular pattern was present (β = 0.346, p < 0.001, probe = 1, pattern = 0, see Supplementary Table **5**). Therefore, this suggests that the animals detect the transition quicker, and in a pattern length dependent manner, when additionally having the repeating pattern in the tone sequence.

### 3.3 Ferrets demonstrate sensitivity to the number of unique elements within a pattern

To further understand what cues are utilised by trained ferrets, we analysed our data according to the number of unique elements within a sequence. We reasoned that if ferrets combine information about spectral statistics with regularity detection, then patterns with more unique frequency elements might be harder to detect than patterns in which one or more frequencies repeated. Figure 5 plots performance by the number of unique frequencies for all pattern lengths; it is clear that performance decreases as the number of unique frequencies within the pattern increases, even within the same pattern length. A mixed effects logistic regression confirmed this observation yielding a negative coefficient for the effect of the number of frequencies on performance (β = −0.366, p < 0.001; see Supplementary Table **6**), as well as main effects of random alphabet (β = 0.126, p < 0.001) and pattern length (β = −0.0843, p < 0.001). These results suggest that ferrets’ ability to detect regularity is constrained by spectral complexity, with reduced performance for sequences containing a greater number of unique frequency elements in addition to the pattern length, suggesting that information on spectral statistics are integrated with regularity detection.

**Figure 5:**
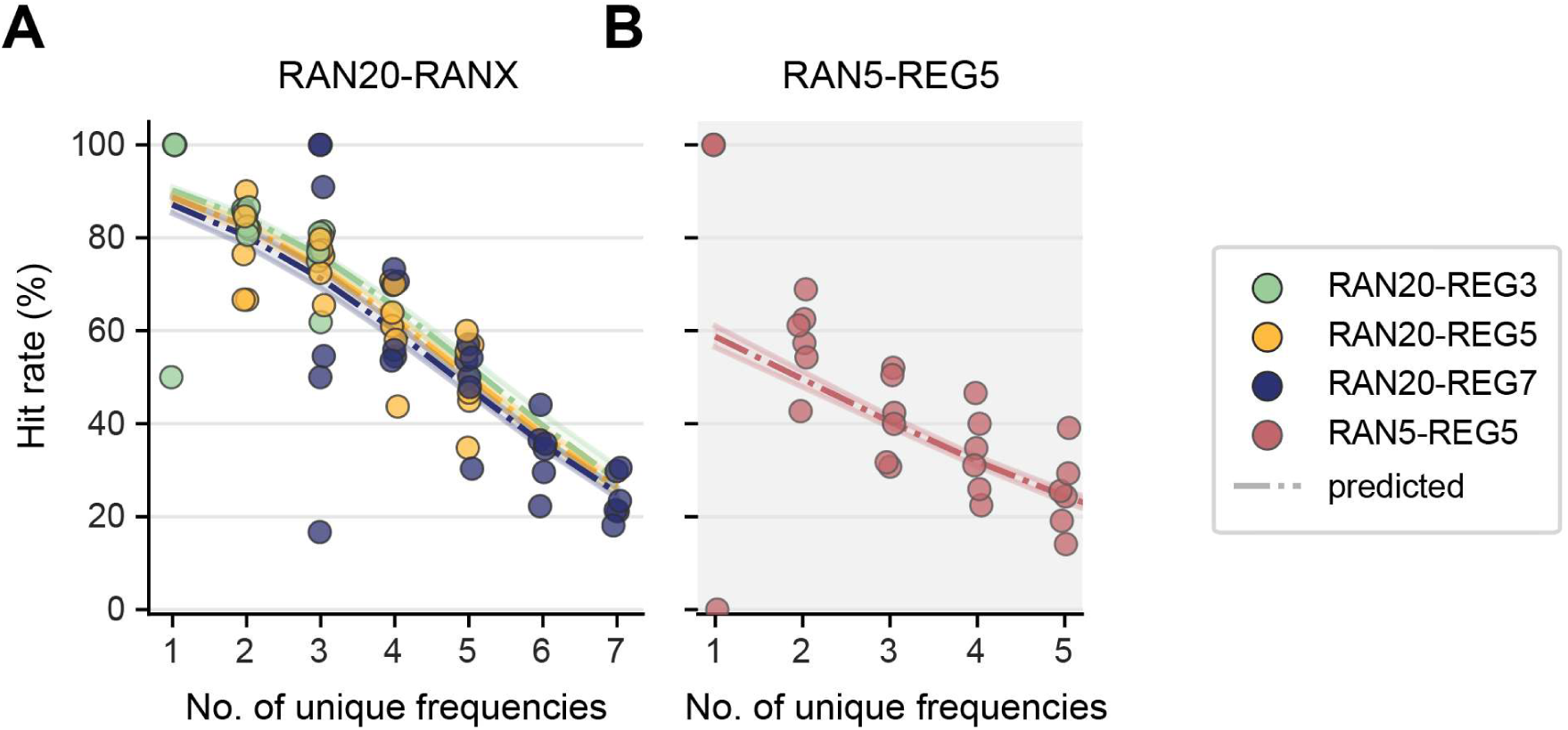
Hit rate for patterns with various unique frequencies. A) Hit rate for each ferret as a function of the number of unique frequencies in the pattern, coloured by pattern length (RAN20-REGX) for RAN20 sequences and B) RAN5 sequences. Dashed lines; predicted values from the mixed effects logistic regression (±95% confident intervals).

### 3.4 Probing ferret behaviour with PPM and PPM-Decay models

Overall, the findings indicate that ferrets are adept at detecting statistical regularities in sound. While they were able to detect regularity for longer patterns, and for trials in which the frequency statistics did not change, performance was worse for longer patterns, and sequences comprised of more unique frequencies. The observed effect of the number of unique frequencies could reflect that ferrets can leverage changes in spectral probability when available. Alternatively (or additionally), this may suggest a limitation of working memory, with patterns that have more unique frequencies placing a greater burden on memory resources. To gain further insight into what factors may shape performance we turned to modelling, using the PPM model, which has been successfully applied to simulate an ideal observer for such stimuli (Barascud et al., 2016, see methods). This model measures the information content of the upcoming tone based on previous tone presentations and context. Information content, measured in bits, quantifies how surprising or unpredictable a stimulus is given its context. One bit corresponds to the information needed to choose between two equally likely outcomes (e.g. two tones). If 20 tones were equally probable, the maximum information content would be log₂(20) ≈ 4.32 bits. However, when an upcoming tone strongly violates the model’s context-based predictions, the information content can exceed what would be expected from frequency alone. In contrast, highly predictable sequences yield values approaching zero. We assume that listeners may detect regularities by identifying a sharp drop in information content, with larger and more abrupt reductions facilitating easier and quicker detection.

As shown in Figure 6A, when applying the PPM model to sequences with a transition from random to regular, information content drops sharply within approximately 3 tones following the transition time (the start of the second repetition). This is in comparison to the consistently high level of information content in the random sequences (grey). This effect is consistent across REG3, REG5 and REG7, conditions. However, in the RAN5-REG5 condition, information content for the random sequence is lower due to the reduced set of possible frequencies. Consequently, the separation between random and regular trials is smaller, and the drop in information content is slightly more delayed. This suggests that under ideal conditions, the model can detect the transition equally well for RAN20-REG3, 5 and 7, but may be delayed for RAN5-REG5. When applied to probe trials (RAN20-RANX), which only differ in the number of frequencies at the transition and without regularity, the model shows a small drop in information content at the transition point (see Figure 6B). However, this drop is very variable, and overlaps with variation in information within random sequences, particularly for larger frequency pools. Together, this suggests an ideal observer would not reliably detect a transition in these conditions.

**Figure 6:**
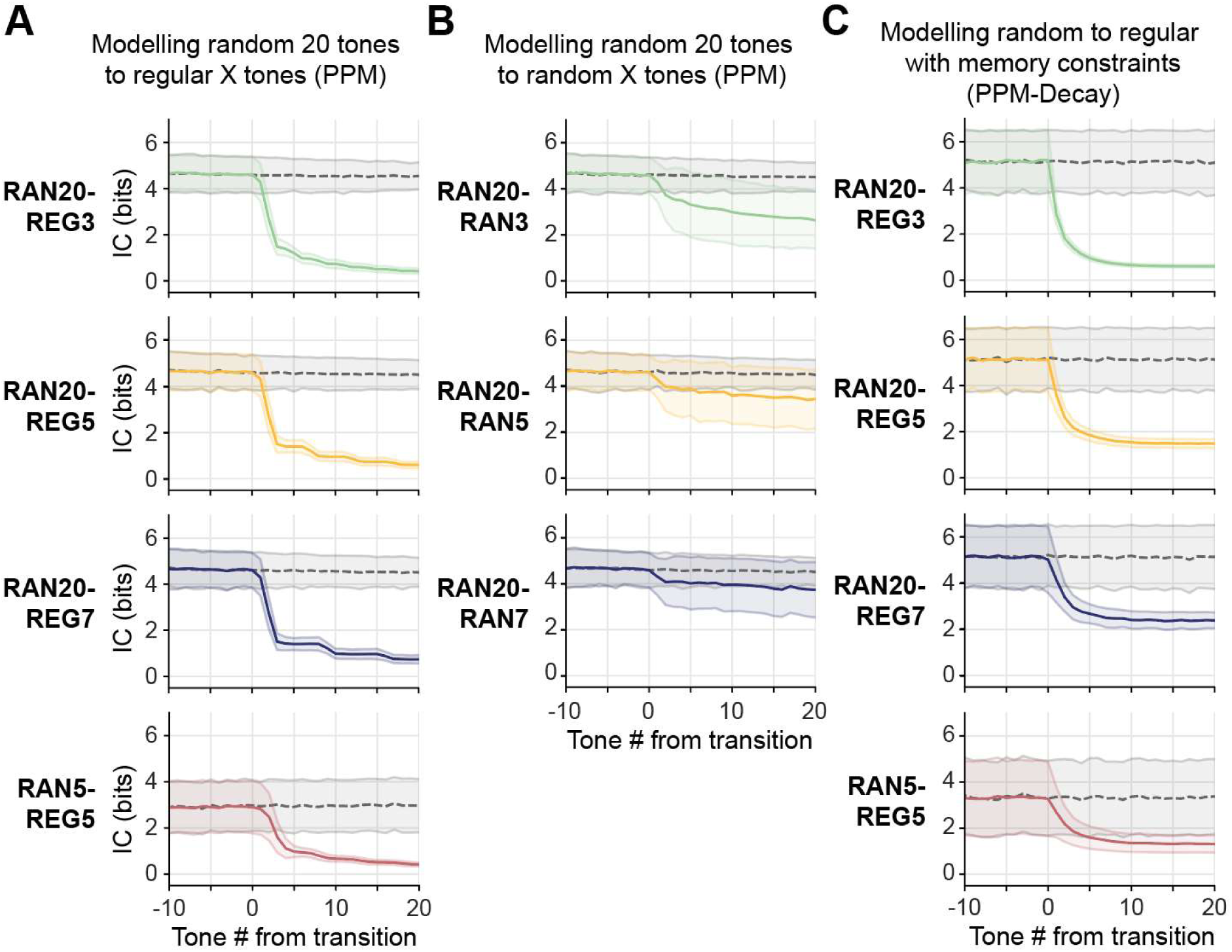
PPM and PPM-Decay model output of tone sequences. A) Average information content (IC; based on the PPM model) for each condition for the RANX-REGX (coloured) and RANX (grey) sequence. B) Average information content (IC; based on the PPM model) for each condition for the RAN20-RANX (coloured) and RAN20 (grey) sequence. C) Average information content (based on the PPM-Decay model with a long-term memory half-life of 0.09) for each condition for the RANX-REGX (coloured) and RANX (grey) sequence. PPM-decay model predictions, with reduced memory capacity. This qualitatively reproduces the impaired performance shown by ferrets for RAN20-REG7 and RAN5-REG5 conditions. Solid/dashed lines are the mean information content across trials for RAN-REG/RAN and RAN trials respectively, shaded regions indicate standard deviation.

Similar to human behavioural studies (Barascud et al., 2016), ferret performance declines as pattern length increases and reaction times increase with longer patterns, although in humans reaction times typically only begin to lag for patterns of above 10 tones than compared to the pattern lengths of 5 and 7 observed in this study. According to the PPM model, an ideal observer with perfect short- and long-term memory should detect the random to regular transition within 3 tones for the unmatched alphabet conditions (RAN20 to REG3, 5 and 7), despite the change in pattern length, and in 5 tones for the matched alphabet condition (RAN5-REG5). However, using a modified version of the model, the PPM-Decay model, which incorporates variable short- and long-term memory constraints, previous research has replicated the delayed detection observed with longer pattern lengths in humans (Harrison et al., 2020). Considering that ferrets, like humans, are unlikely to behave as ideal observers, we explored whether the PPM-Decay model could account for the delays in reaction time with the increases in pattern length for the much shorter patterns used for the ferret behaviour. By adjusting the long-term memory phase half-life in the PPM-Decay model, we limited how long past events are retained in memory. Information content calculations with and without these constraints revealed two key effects (see Figure 6C). First, memory constraints increased the variance of information content and reduced the separation between random and regular conditions as the pattern length increased and is particularly evident in the RAN5-REG5 condition, potentially lowering detection accuracy. The model also shows lower variance than the PPM model in the RAN20-REG3 condition, likely due to reduced memory constraining context and consistently lowering information content. Second, the slope of the information content drop also became shallower with increasing pattern length, and again in the RAN5-REG5 condition, suggesting that more tones are needed to detect the transition.

When comparing the model simulations of information content to ferret behaviour, the PPM-Decay model captures key features of behavioural performance that do not necessarily align with that of an ideal observer model (PPM) or a human listener. Specifically, increased variance in information content was observed for longer patterns and for conditions where the random and regular sequences were drawn from the same frequency set (e.g., RAN5-REG5), resulting in reduced separation between random and regular sequences. This can be seen as proxy for reduced detection accuracy, which is what we observed behaviourally in the ferret with lower detection rates as the pattern length increased and for the RAN5-REG5 condition. For these sequences, the model also showed a delay in the drop of information content following the transition point as pattern length increased, consistent with the longer reaction times observed in the ferrets. These findings suggest that the ferrets’ performance may be shaped by comparable memory constraints, and that the PPM-Decay model offers a plausible computational account of the factors influencing detection under varying pattern complexities.

## 4. DISCUSSION

Here we demonstrate that that, like humans, ferrets can detect the emergence of a novel repeating pattern from a preceding random tone sequence, with performance declining as pattern complexity increases. These findings suggest that the ability to detect novel repeating patterns a fundamental property of the mammalian auditory system. Moreover, these results validate the ferret as a suitable model for behavioural and neural studies of acoustic regularity detection.

We trained a total of six ferrets on a GO/NO-GO task to detect a transition from a random tone sequence to a regular tone sequence. We demonstrated that all ferrets tested could successfully detect the emergence of patterns from three tones in length up to seven tones. This ability to detect novel deterministic patterns has not yet been demonstrated in the ferret until now and provides a unique avenue to investigate parametrically how the auditory system deal with the discovery and perception of auditory patterns. Changing the length of the pattern offers the opportunity to understand how responses change with increasing complexity of the regularity. We observed that detection (hit-rate) decreases from approximately 80% at RAN20-REG3 to 35% at RAN20-REG7 whilst the false alarm rate remained fairly consistent across conditions. Reaction times also increased by just under 200 ms, overall showing a systematic decrease in performance as you increase the pattern length. This pattern of results is consistent with findings in macaques, where reaction times increased and hit-rates decreased as the regularity or coherence of the figure-ground stimuli decreased (Schneider et al., 2018).

Previous work in human listeners showed that they performed similarly to an ideal observer for patterns length under 15 tones (Barascud et al., 2016), detecting the pattern within four tones of the repeated pattern. However, as the pattern length increased to 20 tones, detection required five or more tones. To estimate how many tones are required to detect a pattern requires a method to estimate the motor response time and separate this from the reaction time. This baseline correction is typically achieved through detecting a simple change in frequency of a repeating tone (Barascud et al., 2016) but our ferret subjects would not reliably do this. While we were unable to baseline-correct the ferrets’ reaction times and therefore cannot translate reaction time to a number of tones, it is evident that they do not act as an ideal observer for the pattern lengths presented in this study, as their reaction times increased with increasing pattern duration. Nonetheless, the consistent changes in ferret performance and reaction times with increasing pattern length demonstrate their sensitivity to sequence complexity.

Our findings suggest that ferrets rely on similar mechanisms as humans to detect patterns but may be more limited in certain aspects of regularity detection. We propose that this limitation could stem from memory constraints. Although no experimental data have directly examined the role of memory in pattern detection, very recent work has shown that a listeners explicit short-term memory ability is predictive of the separation of the neural response in EEG between the random and regular sequences (Hu, 2024). Furthermore, Bianco et al. (2023) found that older human listeners detected novel patterns more slowly. This effect was successfully modelled using a PPM-Decay framework with memory constraints, in contrast to an ideal observer with no such limitations. Applying this same model to ferret behaviour, we observed a similar trend in which memory of preceding tones degrades quickly and the drop in information content occurs later in the tone sequence with increasing pattern length. These changes could explain both their decreased performance at longer pattern lengths, and account for their delayed reaction times, if detection relied on a drop in information content below a static threshold. This supports the suggestion that memory limitations may underlie the differences in pattern detection between ferrets and humans. Exploring how these constraints vary across species is a promising direction for future research.

When detecting repeating patterns in these stimuli listeners may rely on auditory cues beyond the sequence of tones that forms the pattern. In our stimuli, one such cue is the change in frequency statistics; specifically, the increased probability of the tones that form the repeating pattern which can facilitate regularity detection despite memory constraints. Recent research has shown that human participants detected these changes most effectively when both the mean and variance of the new tone sequence differed from those of the preceding sequence (Zhao et al., 2025). To test the ferrets’ ability to detect such changes, we presented probe trials which only contained a change in stimulus frequency statistics (i.e. a drop in random frequencies from 20 to 3, 5 or 7) but no repeating patterns. We observed that the ferrets were significantly less likely to respond to the probe in comparison to the regular patterns. Furthermore, we also presented stimuli that contained the same frequencies in the random and regular sequence (RAN5-REG5). All ferrets successfully detected the pattern above chance; however, their performance was lower than in the RAN20-REG5 condition and comparable to the RAN20-REG7 condition. Together, these findings suggest that ferrets accurately detect the emergence of regularity but can benefit from the change in stimulus statistics as an additional cue.

An intriguing observation in our behavioural data is the decline in regularity detection performance as the number of unique frequencies increases. This suggests that greater spectral diversity within the pattern imposes additional cognitive demands, potentially making it harder for ferrets to track multiple frequency bands simultaneously. Such demands may arise because the auditory system processes sound through parallel frequency channels (Bitterman et al., 2008; Chi et al., 2005; Santoro et al., 2014), and increased spectral complexity may strain these mechanisms. This aligns with the idea that detection performance may be influenced not only by memory limitations but also by how spectral information is integrated across channels. While the model used in this study does not account for spectral structure, recent work has shown that models incorporating frequency-band tracking may aid in explaining the neural responses to regular sequences (Zhao et al., 2025) and may offer a more accurate framework for capturing these dynamics.

The observation that patterns with fewer unique frequencies are more readily detected may also hint at an additional detection strategy of rhythm-based detection. Sequences with fewer unique frequencies might give rise to a stronger sense of temporal regularity, similar to detecting the recurring ‘A’ in a pattern like ABACA, which could facilitate performance by engaging rhythm-sensitive mechanisms. This interpretation aligns with a proposal that perhaps animal models may rely more on rhythm to detect changes in an acoustic stimulus (Bianco et al., 2024). Although periodicity was not directly assessed in this study, work has suggests that periodicity could serve as a powerful cue for regularity detection (Asokan et al., 2021; Bianco et al., 2024). Whether the brain encodes the full sequence structure or merely a subset of salient, rhythmically organised elements, and how this may differ between species remains an open question.

In summary, our findings establish the ferret as a valuable non-primate animal model for studying auditory pattern detection. Ferrets can detect the emergence of regularity in sound sequences, with performance shaped by both memory demands and spectral complexity. While they do not behave as ideal observers, their systematic sensitivity to pattern length and frequency structure mirrors key aspects of human auditory behaviour. This positions the ferret as a powerful model for investigating the computational and biological mechanisms of regularity detection. Future work combining behavioural paradigms with neurophysiological recordings will be crucial for uncovering how the ferret auditory system encodes structure in sound, offering comparative insights into the neural basis of auditory cognition across species and bridging the gap between animal models and human neuroimaging findings.

## ACKNOWLEDGEMENTS

This work was supported in whole, or in part, by Wellcome Trust/Royal Society Sir Henry Dale Fellowship Grant 098418/Z/12/A to J.K.B.; European Research Council Consolidator award 771550 SOUNDSCENE to J.K.B; and BBSRC grant BB/P003745/1 to M.C. We would like to thank Eleanor Jones and Zuzanna Slonina, and the rest of the Bizley team for help with behavioural testing of the ferrets.

## 5. SUPPLEMENTARY MATERIAL

**Table 2:** Estimates of each fixed effect in the binomial mixed effects regression model on performance. R^2^ = 0.529; Df = 5849; random effect std. = 0.121.

| Fixed effects | Estimate | Standard error | T | p value | CI 95% |  |
| --- | --- | --- | --- | --- | --- | --- |
|  |  |  |  |  | Lower | Upper |
| **Intercept** | 1.585 | 0.0554 | 28.622 | **< 0.001** | 1.476 | 1.693 |
| **Pattern length** | -0.271 | 0.00423 | -63.911 | **< 0.001** | -0.278 | -0.262 |
| **Random alphabet** | 0.0406 | 0.000765 | 53.114 | **< 0.001** | 0.0391 | 0.0421 |

**Table 3:** Estimates of each fixed effect in the linear mixed effects regression model on reaction time. R^2^ = 0.0993; Df = 5107; random effect std. = 0.0514.

| Fixed effects | Estimate | Standard error | T | p value | CI 95% |  |
| --- | --- | --- | --- | --- | --- | --- |
|  |  |  |  |  | Lower | Upper |
| **Intercept** | 0.626 | 0.0284 | 21.428 | **< 0.001** | 0.554 | 0.665 |
| **Pattern length** | 0.0461 | 0.00238 | 19.366 | **< 0.001** | 0.0414 | 0.0508 |
| **Random alphabet** | 0.00503 | 0.000678 | 6.554 | **< 0.001** | 0.00353 | 0.00654 |

**Table 4:** Model output for mixed effects binomial regression on the p(go) after the transition on probe and pattern trials R^2^ = 0.932; Df = 26; random effect std. = 6.203×10^-10^. Separate model for probe (R^2^ = 0.384; Df = 13; random effect std. = 0.0844) and pattern (R^2^ = 0.964; Df = 13; random effect std. = 0.135) underneath.

| Fixed effects | Estimate | Standard error | T | p value | CI 95% |  |
| --- | --- | --- | --- | --- | --- | --- |
|  |  |  |  |  | Lower | Upper |
| **Intercept** | 2.412 | 0.111 | 21.818 | **< 0.001** | 2.184 | 2.639 |
| **Probe vs pattern (ref = pattern)** | -2.200 | 0.257 | -8.561 | **< 0.001** | -2.727 | -1.671 |
| **Pattern length** | -0.331 | 0.0194 | -17.112 | **< 0.001** | -0.371 | -0.295 |
| **Pattern length × probe** | 0.177 | 0.049 | 3.616 | **0.00126** | 0.0765 | 0.278 |
| Probe |  |  |  |  |  |  |
| **Intercept** | 0.212 | 0.235 | 0.900 | 0.385 | -0.296 | 0.719 |
| **Pattern length** | -0.154 | 0.045 | -3.417 | **0.00459** | -0.251 | -0.0566 |
| Pattern |  |  |  |  |  |  |
| **Intercept** | 2.397 | 0.126 | 18.977 | **< 0.001** | 2.125 | 2.670 |
| **Pattern length** | -0.334 | 0.0194 | -17.213 | **< 0.001** | -0.376 | -0.292 |

**Table 5:** Model output for mixed effects linear regression on reaction time on probe and pattern trials. R^2^ = 0.0704; Df = 3768; random effects std. = 0.0512.

| Fixed effects | Estimate | Standard error | T | p value | CI 95% |  |
| --- | --- | --- | --- | --- | --- | --- |
|  |  |  |  |  | Lower | Upper |
| **Intercept** | 0.842 | 0.0339 | 234.818 | **< 0.001** | 0.775 | 0.908 |
| **Pattern length** | -0.000618 | 0.00479 | 0.129 | 0.861 | -0.00877 | 0.0100 |
| **Probe vs pattern (ref = pattern)** | 0.346 | 0.0831 | 4.167 | **< 0.001** | 0.183 | 0.509 |
| **Pattern length × probe** | 0.0181 | 0.0165 | 1.108 | 0.273 | -0.0143 | 0.0504 |

**Table 6:** Model output for mixed effects logistic regression on performance with regards to unique frequencies within the pattern. R^2^ = 0.186; Df = 104005; random effect std. = 0.311.

| Fixed effects | Estimate | Standard error | T | p value | CI 95% |  |
| --- | --- | --- | --- | --- | --- | --- |
|  |  |  |  |  | Lower | Upper |
| **Intercept** | 0.532 | 0.193 | 2.748 | **0.00560** | 0.152 | 0.911 |
| **Pattern length (PL)** | -0.0843 | 0.0220 | -3.834 | **< 0.001** | -0.127 | -0.412 |
| **Random alphabet (RA)** | 0.126 | 0.00583 | 21.59 | **< 0.001** | 0.114 | 0.137 |
| **Unique frequencies** | -0.366 | 0.0319 | -11.458 | **< 0.001** | -0.428 | -0.303 |
| **PL × Unique freqs** | 0.00695 | 0.00456 | 1.522 | 0.128 | -0.00200 | 0.0158 |
| **RA × Unique freqs** | -0.00967 | 0.00146 | -6.634 | **< 0.001** | -0.0125 | -0.00681 |

**Table 7:** Model output for mixed effects linear regression on reaction time with regards to unique frequencies within the pattern. R^2^ = 0.0252; Df = 56525; random effect std. = 0.0394. RAN20-REG3: R^2^ = 0.0217; Df = 25462; random effect std. = 0.0655. RAN20-REG5: R^2^ = 0.0217; Df = 15857; random effect std. = 0.0280. RAN20-REG7: R^2^ = 0.0113; Df = 5618; random effect std. = 0.0485. RAN5-REG5: R^2^ = 0.0034; Df = 9586; random effect std. = 0.0274.

| Fixed effects | Estimate | Standard error | T | p value | CI 95% |  |
| --- | --- | --- | --- | --- | --- | --- |
|  |  |  |  |  | Lower | Upper |
| **Intercept** | 0.686 | 0.0456 | 15.038 | **< 0.001** | 0.597 | 0.775 |
| **Pattern length (PL)** | 0.0530 | 0.00604 | 8.770 | **< 0.001** | 0.0412 | 0.0649 |
| **Random alphabet (RA)** | -0.0123 | 0.00169 | -7.290 | **< 0.001** | -0.0157 | -0.00903 |
| **Unique frequencies** | 0.0527 | 0.00999 | 5.279 | **< 0.001** | 0.0332 | 0.0723 |
| **PL × Unique freqs** | -0.0156 | 0.00130 | -12.026 | **< 0.001** | -0.0182 | -0.0131 |
| **RA × Unique freqs** | 0.00418 | 0.000442 | 9.450 | **< 0.001** | 0.00331 | 0.00505 |
| RAN20-REG3 |  |  |  |  |  |  |
| **Intercept** | 0.545 | 0.0340 | 16.03 | **< 0.001** | 0.479 | 0.612 |
| **Unique frequencies** | 0.110 | 0.00731 | 15.044 | **< 0.001** | 0.0956 | 0.124 |
| RAN20-REG5 |  |  |  |  |  |  |
| **Intercept** | 0.729 | 0.0265 | 27.529 | **< 0.001** | 0.677 | 0.781 |
| **Unique frequencies** | 0.0523 | 0.00523 | 9.987 | **< 0.001** | 0.0420 | 0.0625 |
| RAN20-REG7 |  |  |  |  |  |  |
| **Intercept** | 0.835 | 0.0440 | 18.966 | **< 0.001** | 0.748 | 0.921 |
| **Unique frequencies** | 0.0229 | 0.00659 | 3.481 | **< 0.001** | 0.0100 | 0.0358 |
| RAN5-REG5 |  |  |  |  |  |  |
| **Intercept** | 0.881 | 0.0237 | 37.108 | **< 0.001** | 0.835 | 0.928 |
| **Unique frequencies** | -0.00448 | 0.00597 | -0.750 | 0.453 | -0.0162 | 0.00722 |

